# The cost of correcting for error during sensorimotor adaptation

**DOI:** 10.1101/2021.01.13.426535

**Authors:** Ehsan Sedaghat-Nejad, Reza Shadmehr

**Affiliations:** Laboratory for Computational Motor Control, Department of Biomedical Engineering, Johns Hopkins School of Medicine, Baltimore MD

**Author notes:** **Correspondence:** Ehsan Sedaghat-Nejad or Reza Shadmehr, Traylor Building Room 416, Johns Hopkins School of Medicine, 720 Rutland Ave., Baltimore, MD 21205. or.

## Abstract

Learning from error is often a slow process. To accelerate learning, previous motor adaptation studies have focused on explicit factors such as reward or punishment, but the results have been inconsistent. Here, we considered the idea that a movement error carries an implicit cost for the organism because the act of correcting for error consumes time and energy. If this implicit cost could be modulated, it may robustly alter how the brain learns from error. To vary the cost of error, we considered a simple saccade adaptation task but combined it with motion discrimination: movement errors resulted in corrective saccades, but those corrections took time away from acquiring information in the discrimination task. We then modulated error cost using coherence of the discrimination task and found that when error cost was large, pupil diameter increased, and the brain learned more from error. However, when error cost was small, the pupil constricted, and the brain learned less from the same error. Thus, during sensorimotor adaptation, the act of correcting for error carried an implicit cost for the brain. Modulating this cost affects how the brain learns from error.

## Introduction

In machine learning, the error in the output of a network is evaluated by a loss function that depends on the difference between the output of the network, and the desired one. This loss function is a mathematical description of the cost of error, which in turn is the principal driver of how much the network should learn from error. In analogy to machine learning, during sensorimotor tasks learning in humans also depends on a loss function (Kording and Wolpert, 2004) that tends to grow with error magnitude (Marko et al., 2012). This implies that in principle, altering the landscape of the sensorimotor loss function should affect the rate of learning. However, it has been difficult to find ways to modulate the sensorimotor loss function.

Previous approaches have considered inducements such as monetary reward or punishment, thus associating an explicit cost to the movement error. Initial results suggested that associating error magnitude with monetary loss was effective in accelerating adaptation (Galea et al., 2015; Song and Smiley-Oyen, 2017). However, later work questioned this observation (Quattrocchi et al., 2018). Other work noted that associating error magnitude with reward enhanced the rate of adaptation (Nikooyan and Ahmed, 2015), but that result was also difficult to replicate (Galea et al., 2015; Spampinato et al., 2019). More recent results suggest that even when there is an effect of reward on rate of learning, it acts primarily through recruitment of the explicit, cognitive component of adaptation, not the implicit, unconscious component (Codol et al., 2018).

Here, we began with the observation that movement errors are often followed by corrective actions. For example, if a saccadic eye movement misses the target, the resulting error encourages the brain to generate a corrective saccade. However, corrective movements carry an implicit cost in that they consume time, which in turn delays acquisition of reward (Shadmehr et al., 2010). Thus, a natural loss function for movement error is the time that is expended in the act of producing the correction. If this time could be linked with a utility, then the landscape of the loss function may be altered, resulting in modulation of learning rates.

To test this idea, we designed a paradigm that combined saccade adaptation with decision-making in a random dot motion discrimination task. Like traditional saccade adaptation tasks, subjects made a saccade toward a visual target and experienced an error that required a corrective movement. However, unlike traditional tasks, the corrective saccade carried a cost: it consumed time needed to acquire information for the decision-making task. We varied this cost via motion coherence of the random dots in the discrimination task and found that modulating the cost of correcting for error was a robust factor in how much the brain learned from error.

## Results

Subjects made center-out horizontal saccades to a visual target (a green dot, 0.5×0.5^o^) at ±15°. At the conclusion of their primary saccade, they were presented with an image that contained random dot motion (Fig. 1A). The objective was to detect the direction of motion of the random dots, which was either upward or downward, and was reported by making a vertical saccade. After this vertical saccade, feedback was provided regarding decision accuracy.

**Fig 1.**
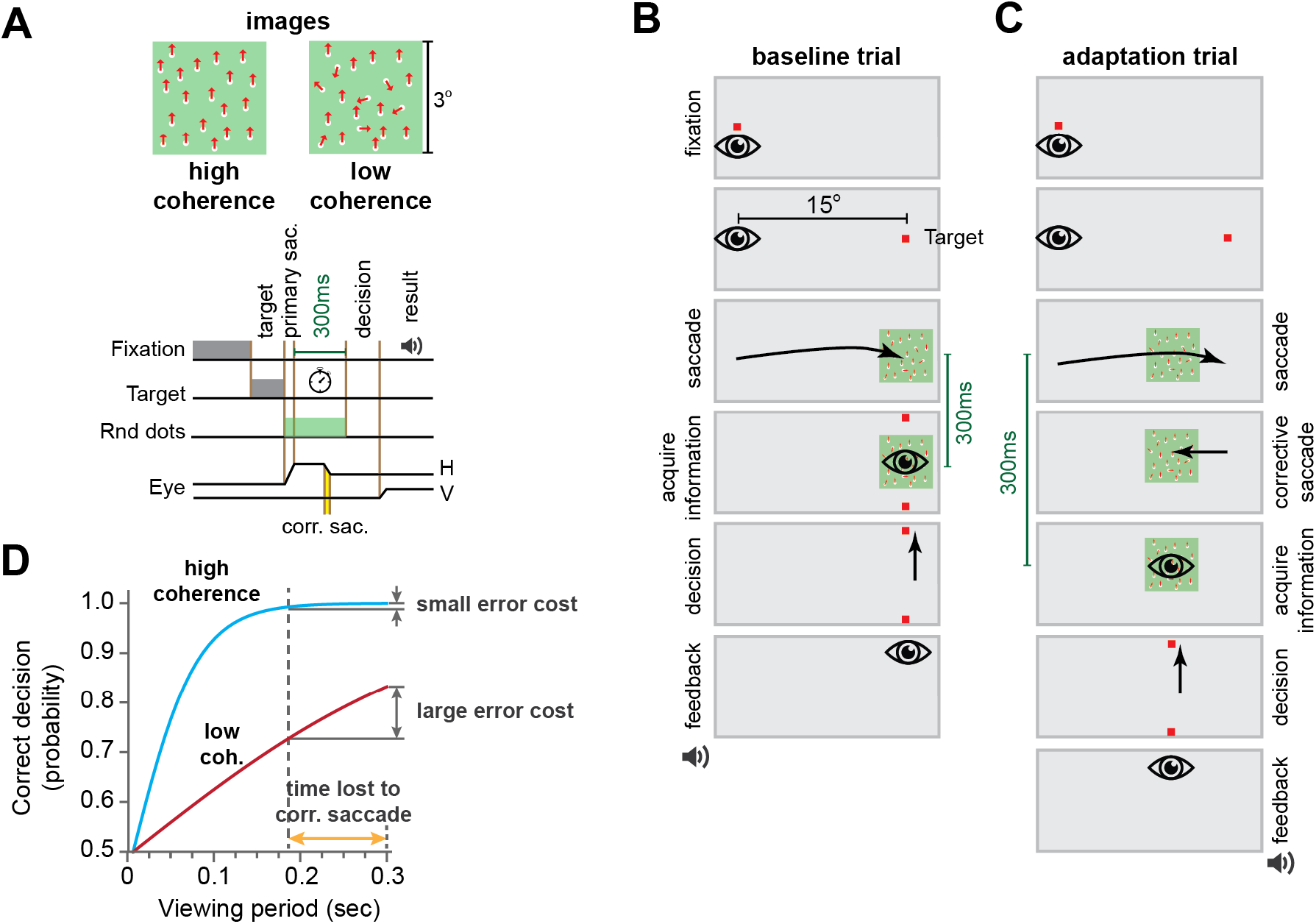
Experiment design. **(A)** Each trial began with a random fixation interval (250-750ms) after which a primary target was placed at 15° randomly to the right or left along the horizontal axis. After conclusion of primary saccade, subjects were presented with a random dot motion image with high or low coherence. The task objective was to detect the direction of motion of the random dots, which was either upward or downward, and was reported by making a vertical saccade. After this vertical saccade, feedback was provided regarding decision accuracy. Subjects had only 300 ms from the end of their primary saccade to view the image. Thus, the time consumed by the corrective saccade reduced the time available to place the image on the fovea. **(B)** During the first 100 trials of each session (baseline block) the random dot motion was presented at the primary target location. **(C)** During adaptation trials the image was consistently placed at a location 5° away from the primary target which acted as a movement error and induced a corrective saccade. **(D)** Hypothetical error cost landscape. The period spent correcting for error carried a cost that depended on stimulus coherence. Low coherence images induced large cost and high coherence stimuli induced a small cost in terms of decision accuracy.

In the baseline block, the image was centered at the target (Fig. 1B). However, during the adaptation block the image was centered 5° away from the target (Fig. 1C). As a result, during adaptation the subjects made a saccade to the target, and then followed that with a corrective saccade to a location near the center of the image (Supplementary Fig. 1). Importantly, the movement error and the resulting corrective saccade carried a cost because the subject had only 300 ms from the end of their primary saccade to view the image. Thus, if the subject learned from movement error and adapted their primary saccade, the corrective saccade would consume less time, allowing them to view the random dots for a longer period, and therefore arrive at a more accurate decision.

To modulate the landscape of the loss function, we varied the coherence of the random dots. We assumed that as the subject viewed the random dots, the brain accumulated evidence for each possibility (upward or downward motion). As shown in Fig. 1D, the evidence accumulation is roughly the temporal integral of the instantaneous difference between the number of dots that moved upward vs. downward, and thus grows faster when the motion is more coherent (more of the dots move in a single direction). As a result, for the low coherence image the time lost in correcting the error has a large effect on decision accuracy. In contrast, for the high coherence image, expending the same period of time has little or no effect on decision accuracy (Pilly and Seitz, 2009). Thus, by varying motion coherence, we varied the cost of error in terms of decision accuracy, which we hypothesized would result in changes in learning rates.

When the target was presented to one side of the screen, motion coherence of the image was always low. As a result, the corrective saccade was costly because it took precious time away from viewing the image. However, when the target was presented to the other side of the screen, motion coherence was high, thus making time expenditure less costly. Critically, probability of success was higher for the stimulus that had high coherence. If reward rate is the principal modulator of adaptation, then adaptation should be faster toward the side that was more rewarding (high coherence). On the other hand, if the error cost is the principal factor, then adaptation should be faster toward the side for which error was more costly (low coherence).

### Cost of error increased both the rate and the asymptote of adaptation

In Exp. 1 (n=20 subjects), following baseline trials (Fig. 1B), subjects made saccades to a target that was always associated with large error cost (low image coherence), and to another target that was always associated with small error cost (high image coherence) (trial structure is shown in Fig. 2A). In baseline trials, as well as during adaptation, the probability of a correct decision was much higher for the small cost target (Fig. 2D, RM-ANOVA, trials 101-650, main effect of cost, F(1,19)=56.831, p<0.0005). Yet, the subjects learned more from errors that carried a large cost (Fig. 2B), as indicated by the fact that adaptation rate was faster for the low coherence image (RM-ANOVA on amplitude change, trials 101-650, main effect of trial F(32,608)=76.099, p<0.0005, and trial by cost interaction F(32,608)=2.515, p<0.0005).

**Fig 2.**
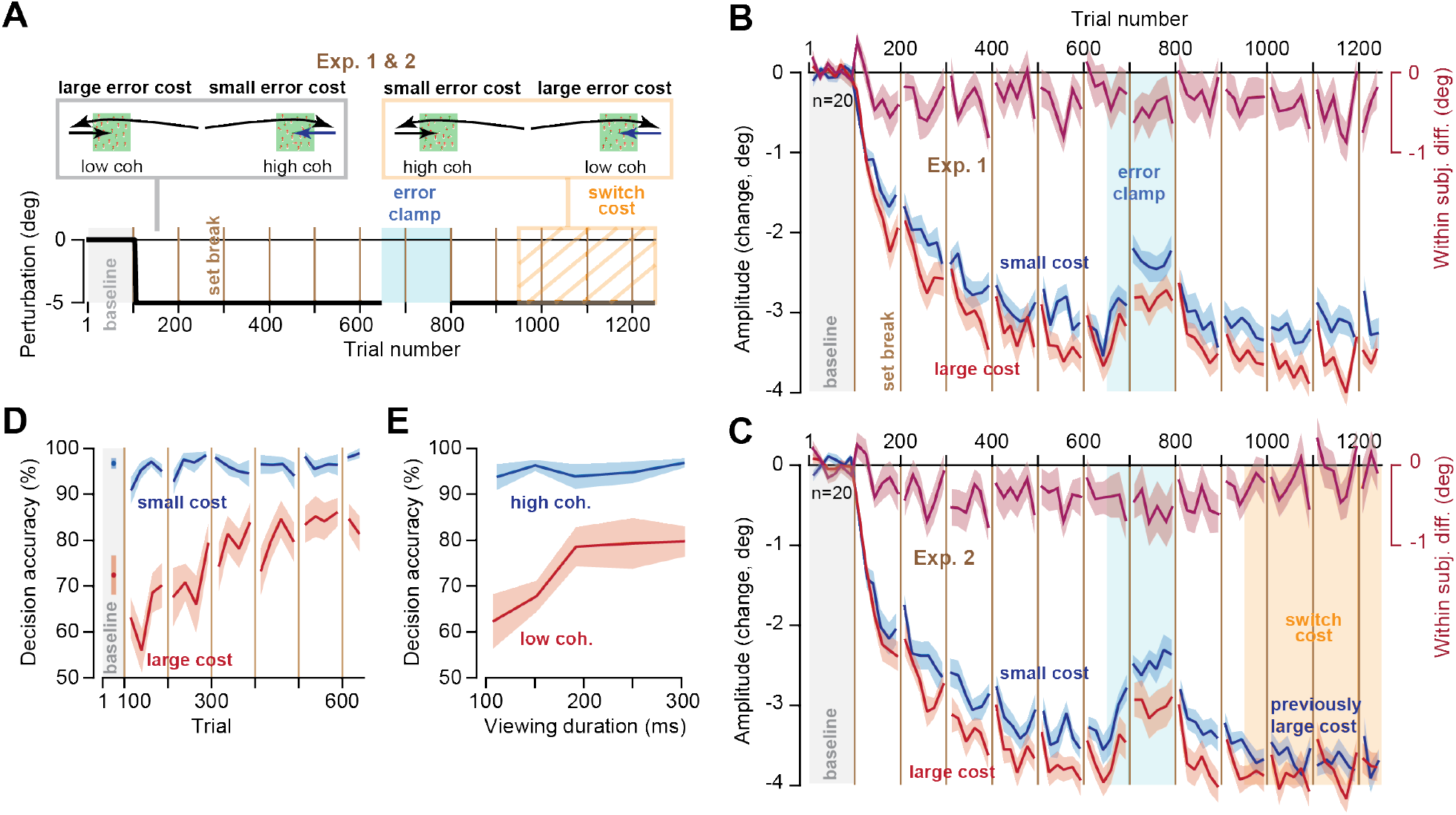
Cost of error modulated rate and asymptote of learning. **(A)** Subjects experienced 550 adaptation trials followed by 150 error clamp trials. In Exp. 1, this was followed by another 450 adaptations trials (trials 801-1250) which established the asymptote of performance. In Exp. 2, after an initial 150 adaptation trials (trials 801-950), at trial 951 (switch cost) suddenly and without warning the side that previously displayed large cost stimuli (low coherence) switched to displaying small cost stimuli (high coherence). **(B & C)** Amplitude change of primary saccades with respect to baseline. Large cost of error increased both the rate and the asymptote of adaptation. **(D)** Decision accuracy for small and large cost stimuli over the course of the experiment. Reward rate was consistently higher for the high coherence (small error cost) image. **(E)** Actual error landscape (compare with hypothetical landscape shown in Fig. 1D). A change in viewing period produced large changes in decision accuracy for the low coherence image, but it had little or no consequence for the high coherence stimulus. Bin size in B & C is 8 trials, in D is 12 trials, and in E is 50 ms. Error bars are SEM.

Following a block of adaptation trials, we imposed a block of error-clamp trials that eliminated movement error. As expected, without errors to sustain adaptation saccade amplitude returned toward baseline levels (Fig. 2B, RM-ANOVA, trials 651-800, main effect of trial, F(8,152)=14.787, p<0.0005, no trial by cost interaction, F(8,152)=0.746, p=0.65). Following the error-clamp block, further training brought performance toward a plateau (Fig. 2B, trials 951-1250). However, adaptation remained higher for the side with the larger error cost: RM-ANOVA, trials 951-1250, main effect of cost F(1,19)=5.439, p=0.031).

Thus, the rate of adaptation was greater toward the target that carried a large cost of error, not the target that carried a greater probability of success. More importantly, the asymptote of adaptation was also greater when the error cost was larger.

### Increasing the cost of error rescued low adaptation

If error cost is a causal mechanism that modulates learning from error, then a change in error cost should produce a change in adaptation. Because Exp. 1 had established that the asymptote of adaptation was greater for the stimulus with large error cost, we checked whether we could rescue adaptation by increasing the error cost.

In Exp. 2, subjects (n=20) began with stimuli that were identical to Exp. 1: large error cost to one side, small error cost to the other. However, at trial 951 (Fig. 2A, switch cost), suddenly and without warning the side that previously displayed large cost images (low coherence) switched to displaying small cost images (high coherence). Similarly, the side that previously displayed small cost images switched to displaying large cost images.

During the initial phase of the experiment (trials 101-650), adaptation rate was faster toward the side that carried a large error cost (Fig. 2C, RM-ANOVA, trial by cost interaction, F(32,608)=2.461, p<0.0005), thus confirming the findings of Exp. 1. Following an error clamp period and initial relearning, we switched the stimuli. This switch from small to large cost coincided with convergence of saccade amplitudes for the two sides (Fig. 2C, RM-ANOVA, trials 801-1250, main effect of trial by cost interaction, F(26,494)=2.781, p<0.0005, no main effect of cost, F(1,19)=1.6, p=0.221). We next compared saccade amplitude during trials 801-1250 in Exp. 1 when there was no switch in cost, with the condition in which the cost switched (Exp. 2). As illustrated in Supplementary Fig. 2, switch in cost appeared to rescue a zero slope learning curve to one that exhibited further learning (Multivariate test, main effect of trial by switch interaction, F(26, 13)=2.522, p=0.042).

It is noteworthy that adaptation rate was greater for the large cost stimulus, despite the fact that the stimulus on the opposite side was more rewarding (Fig. 1D, RM-ANOVA on probability of success, trials 101-650, main effect of cost F(1,19)=56.831, p<0.0005, as well as a trial by cost interaction F(21,399)=4.624, p<0.0005). The consequences of greater reward for the small cost stimulus was readily visible in the reaction time of the primary saccades: as in many previous experiments (Manohar et al., 2015; Milstein and Dorris, 2007; Sedaghat-Nejad et al., 2019b; Shadmehr and Ahmed, 2020; Takikawa et al., 2002; Yoon et al., 2018, 2020), saccades toward the more rewarding stimulus exhibited a shorter reaction time (Supplementary Fig. 3, RM-ANOVA, trials 101-650, Exp. 1, main effect of cost, F(1,19)=9.162, p=0.007, trial by cost interaction, F(32,608)=2.288, p<0.0005). That is, greater reward rate was associated with greater vigor (earlier reaction time), but not greater adaptation. Rather, adaptation rate was higher for the stimulus that carried a greater cost.

Our experiments were based on the assumption that the time period spent correcting for error carried a cost that depended on stimulus coherence (Fig. 1D). To check the validity of this assumption, we quantified the relationship between decision accuracy and viewing time for each stimulus. For the high coherence stimulus, a change in the viewing period produced little or no change in decision accuracy (Fig. 2E). On the other hand, for the low coherence stimulus a change in viewing period produced large changes in decision accuracy (Fig. 2E, interaction of viewing period by decision accuracy, F(4,76)=3.513, p=0.009). This confirmed that the time spent correcting for the movement error carried little or no cost for the high coherence stimulus (small cost), whereas the same expenditure was quite costly for the low coherence stimulus (large cost).

In summary, adaptation rate was greater toward the stimulus that carried a greater error cost, not the stimulus that was more rewarding. When the error cost increased (switch cost), so did the asymptote of performance, suggesting a causal relationship between the cost of error and adaptation.

### Cost of error increased learning from error but not retention

The fact that cost of error modulates the rate of adaptation as well as the asymptote of performance raises the question of whether this cost affects sensitivity to error, trial-to-trial retention, or both. To consider these possibilities, in Exp. 3 we implemented a spontaneous recovery paradigm and then analyzed the results using a state-space model of adaptation (Albert and Shadmehr, 2018).

In Exp. 3 (Fig. 3A), subjects (n=20) began with stimuli that were identical to Exps. 1 and 2: large error cost to one side, small error cost to the other. We again observed that the adaptation rate was faster toward the stimulus with large error cost (Fig. 3B, RM-ANOVA, trials 101-650, trial by cost interaction, F(32,608)=2.944, p<0.0005). After the initial adaptation period we reversed the direction of movement errors (trials 651 to 750) to induce “extinction”, resulting in a sharp change in saccade amplitude toward baseline. Following error reversal subjects experienced a long period of error clamp trials (trials 751-1200). As expected (Ethier et al., 2008), during the error clamp period saccade amplitude exhibited spontaneous recovery toward the adapted state (Fig. 3B, trials 751-800, RM-ANOVA, main effect of trial, F(3,57)=24.709, p<0.0005).

**Fig 3.**
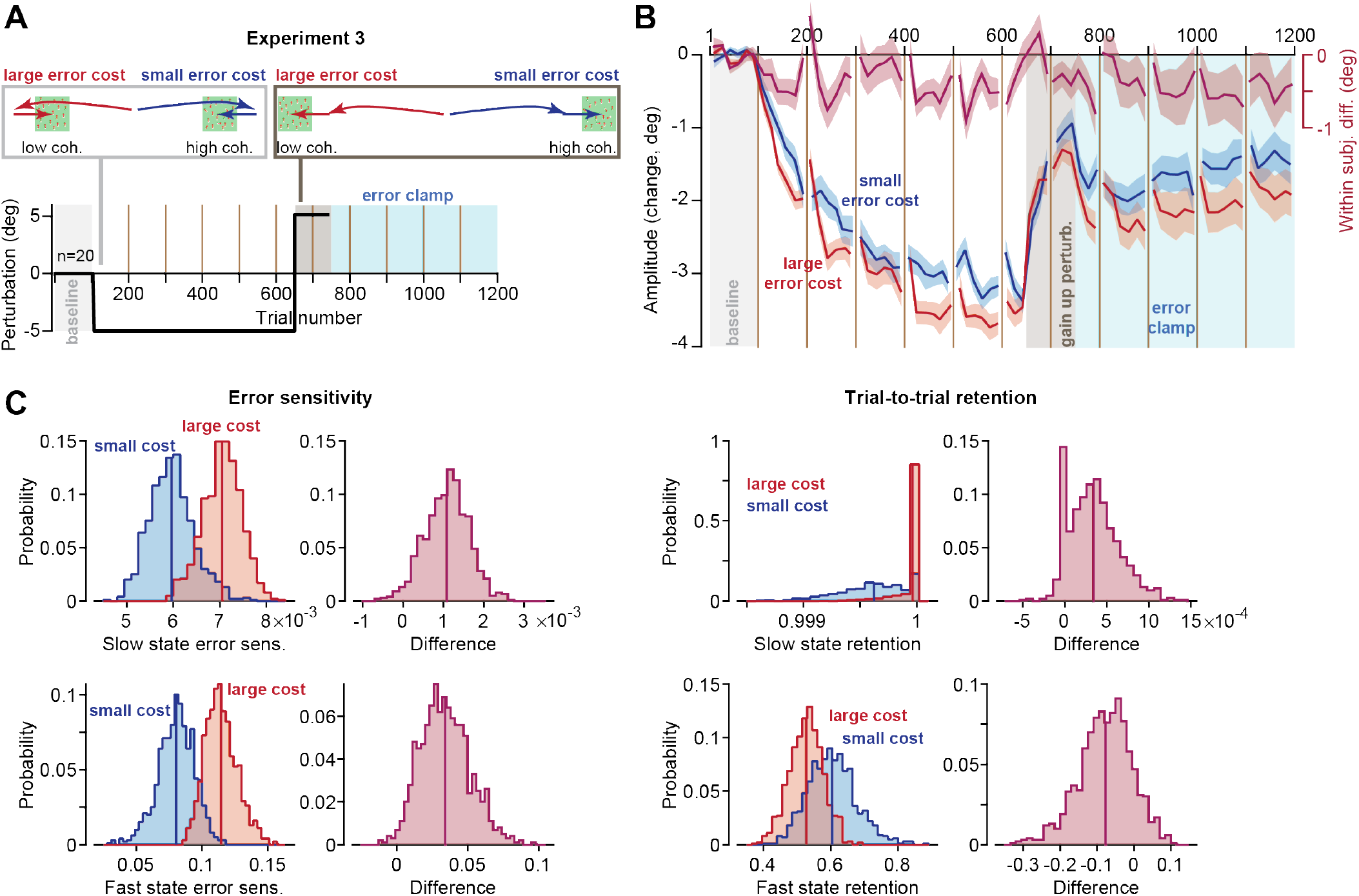
Cost of error affected sensitivity to error but not retention. **(A)** Experiment design of spontaneous recovery paradigm. **(B)** Amplitude change of primary saccades with respect to baseline. **(C)** A two-state model was fitted to the data and error sensitivity and retention were estimated for the fast and slow states using a bootstrap procedure. The plots show the resulting distribution of parameter values and the differences in parameter values due to change in cost. Bin size in B is 8 trials. Error bars are SEM.

We next applied a state space model to estimate error sensitivity and trial-to-trial retention. When the cost of error was large, error sensitivity was elevated for both the slow (Fig. 3C, paired t-test, p=0.019) and the fast state (Fig. 3C, paired t-test, p=0.023). In contrast, cost of error did not appear to affect trial-to-trial retention (Fig. 3C, slow state, paired t-test, p=0.119; fast state, paired t-test, p=0.846).

To check the robustness of this result, we reconsidered the data in Exp. 1, with the caveat being that because this experiment did not contain a spontaneous recovery period, we did not have sufficient power to consider a two-state model, and thus fitted a single-state set of equations. We again found that error sensitivity was larger for the large cost target, with no significant effect on the trial-to-trial retention (small cost and large cost error sensitivity were 14.0×10^−3^ ± 1.95×10^−3^ and 17.5×10^−3^ ± 1.56×10^−3^, within-subject difference was 3.5×10^−3^ ± 1.97×10^−3^, in contrast, we found no significant difference between estimated trial-to-trial retention, small cost and large cost trial-to-trial retentions were 0.9954 ± 7.34×10^−4^ and 0.9961 ± 6.66×10^−4^, within-subject difference was 6.7×10^−4^ ± 9.1×10^−4^, paired t-test, p=0.23).

In summary, cost of error affected adaptation by up-regulating sensitivity to error, not retention.

### Faster adaptation provided more viewing time to perform motion discrimination

We had assumed that adaptation would afford subjects more time to view the image, and thus help them make more accurate decisions. To check for this, we combined the data for the three experiments (n=60 subjects). As expected, large error cost coincided with a faster rate and a greater extent of adaptation (Fig. 4A, RM-ANOVA, trials 101-650, main effect of trial, F(32,1888)=233.864, p<0.0005, main effect of cost, F(1,59)=9.245, p=0.004, and trial by cost interaction, F(32,1888)=3.735, p<0.0005). This increased rate of adaptation for the large cost stimulus provided more time to view the moving dots (Fig. 4B, RM-ANOVA, trials 101-650, main effect of trial, F(32,1888)=84.999, p<0.0005, main effect of cost, F(1,59)=6.841, p=0.011, and trial by cost interaction, F(32,1888)=3.072, p<0.0005). Finally, as saccades adapted and the viewing period increased, so did decision accuracy (Fig. 4C). But as expected, the impact of the increased viewing period on decision accuracy was much greater when time was more valuable (i.e., low coherence stimulus, Fig. 4C, main effect of trial by cost interaction, F(21,1239)=4.994, p<0.0005).

**Figure 4.**
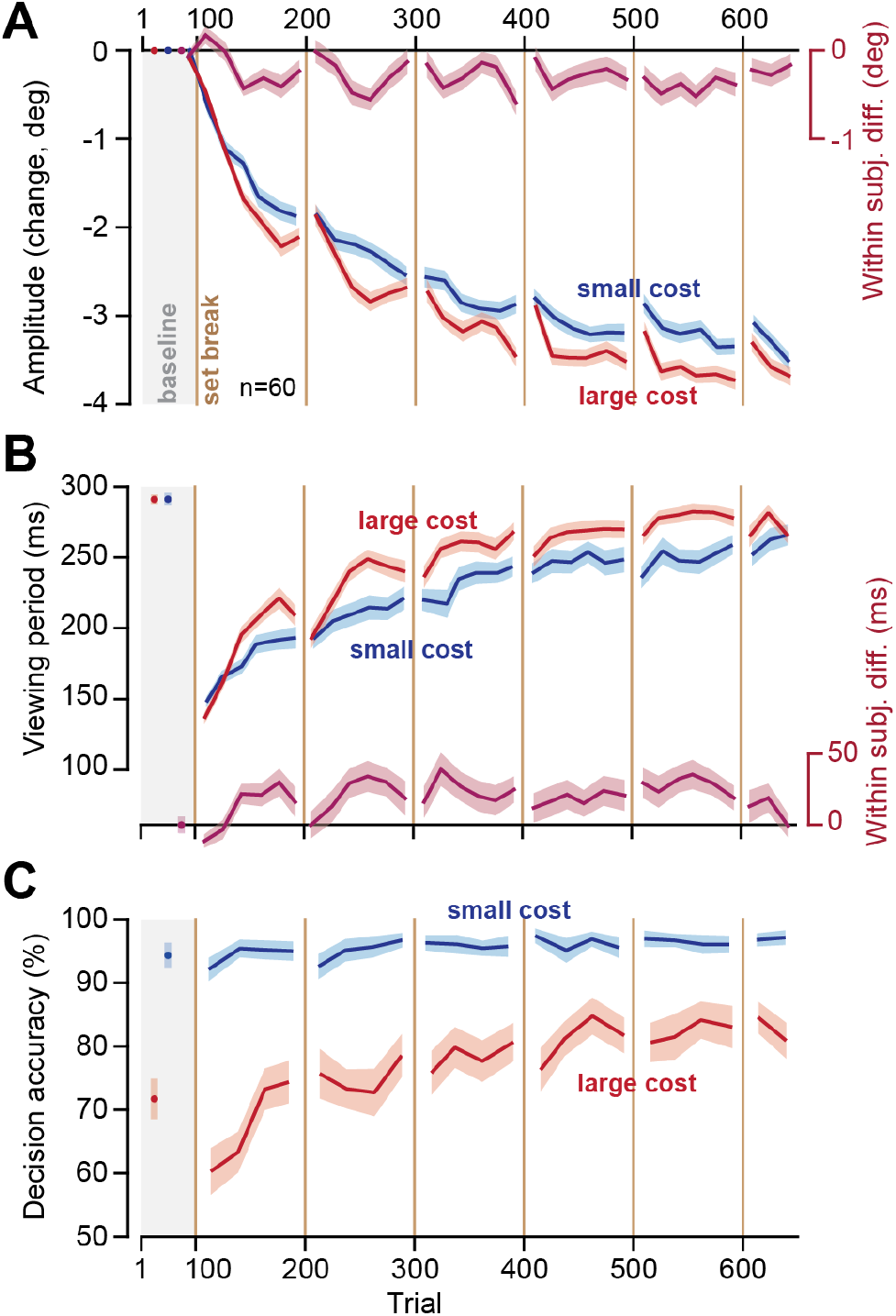
Faster adaptation of the primary saccade provided more viewing time to perform motion discrimination. **(A)** Change in primary saccade amplitude from baseline. **(B)** Image viewing period over the course of adaptation. Faster adaptation provided more viewing time to perform motion discrimination. **(C)** Decision accuracy over the course of the adaptation. The impact of the increased viewing period on decision accuracy was much greater for large cost stimulus as compared to the small cost stimulus. Data are from Exp. 1-3 combined. Bin size in B & C is 8 trials and in C is 12 trials. Error bars are SEM.

### Control experiment: eliminating the error cost equalized rates of adaptation

There is a potential confound in our interpretation: the task was harder for the low coherence stimulus. Thus, it is possible that learning rate was not driven by cost of error, but rather the difficulty of the task. To test for this, we performed a control experiment in which the cost of error was equal for the two stimuli, but task difficulty was greater for one of them (Fig. 5A).

**Fig 5.**
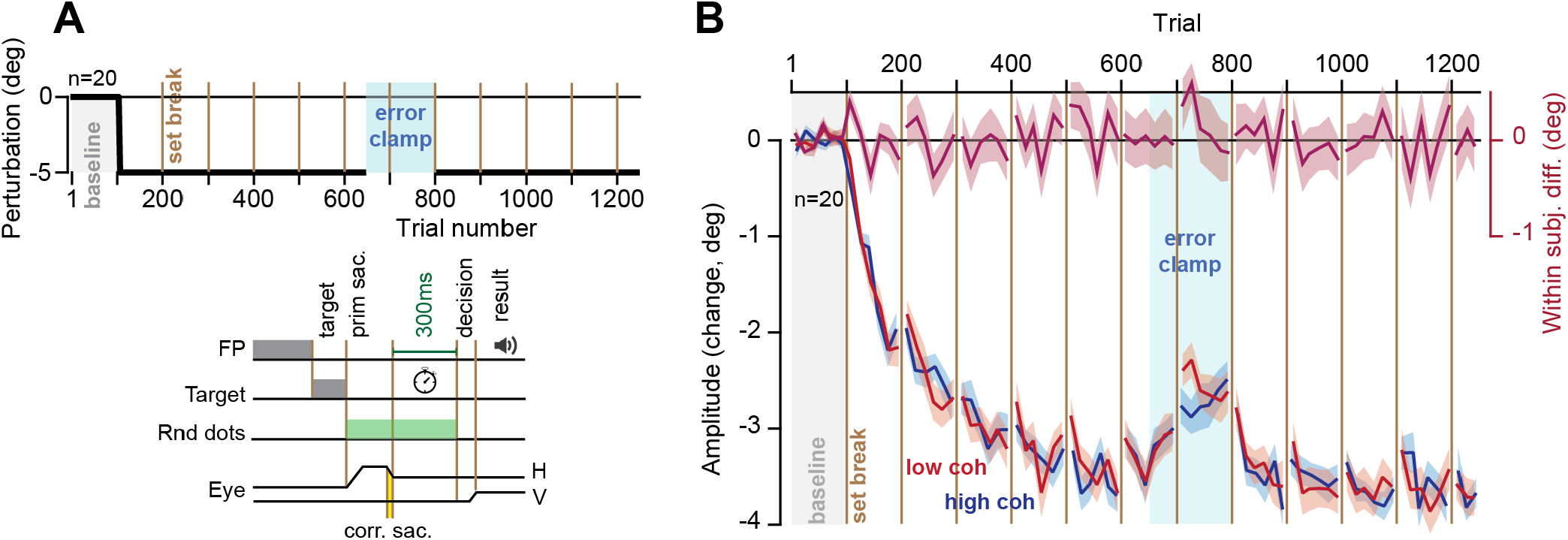
Control experiment: eliminating the error cost equalized rates of adaptation. **(A)** Block design (top panel) was identical to Exp. 1. Trial design was also very similar with one crucial difference; the time to view the random dot image did not start until conclusion of the corrective saccade. This change effectively provided subjects with 300 ms to view the random dots regardless of the primary saccade amplitude. **(B)** Amplitude change in primary saccades with respect to baseline. We found no significant within-subject difference in either rate or asymptote of adaptation between the low and high coherence stimuli. Bin size in B is 8 trials. Error bars are SEM.

As before, subjects (n=20) made a primary saccade to targets at ±15°, and again were presented with an image that was centered 5° away. However, unlike the main experiments, in this control experiment subjects were provided with 300 ms to view the random dot image regardless of the primary saccade amplitude. That is, the time allowed to view the image did not start until conclusion of the corrective saccade (Fig. 5A). With this subtle change we removed the cost associated with the movement error: now the time spent making a corrective saccade did not take away time from viewing the image.

As before, decision accuracy was greater for the side that contained the high coherence image, thus confirming that the task on one side remained more difficult than the other (RM-ANOVA, trials 101-650, main effect of coherence, F(1,19)=60.096, p<0.0005).

However, while saccade amplitude exhibited adaptation (Fig. 5B, RM-ANOVA, trials 101-650, main effect of trial, F(32,608)=74.049, p<0.0005), there were now no significant within-subject differences between the low and high coherence stimuli (Fig. 5B, RM-ANOVA, trials 101-650, no main effect of coherence, F(1,19)=0.018, p=0.895). Furthermore, we found no significant within-subject difference in the asymptotic learning between the two types of stimuli (Fig. 5B, RM-ANOVA, trials 951-1250, no main effect of coherence, F(1,19)=0.008, p=0.931).

In summary, in this control experiment we found that eliminating the cost of error, while maintaining the difference in task difficulty, eliminated modulation of learning rates.

### Pupil dilation coincided with increased cost of error

What might be the neural mechanism that links cost of error with adaptation? To approach this question, we looked at pupil dilation as a proxy for activation of the brainstem neuromodulatory system (Vazey et al., 2018). We measured pupil diameter as subjects fixated the center target and found that at the onset of each block the pupil was dilated, but then progressively constricted as the trials continued (Fig. 6A). Following the set break, the pupil once again became dilated. These patterns were present in both the main experiment and the control experiment (Fig. 6A, RM-ANOVA, trials 101-650, main effect of trial, main experiments: F(1,59)=22.642, p<0.0005, and control experiment: F(1,19)=10.975, p<0.0005). If we view pupil diameter as a proxy for arousal, then it appears that there was a general decline in arousal within each block of trials, followed by sharp recovery due to set breaks.

**Fig 6.**
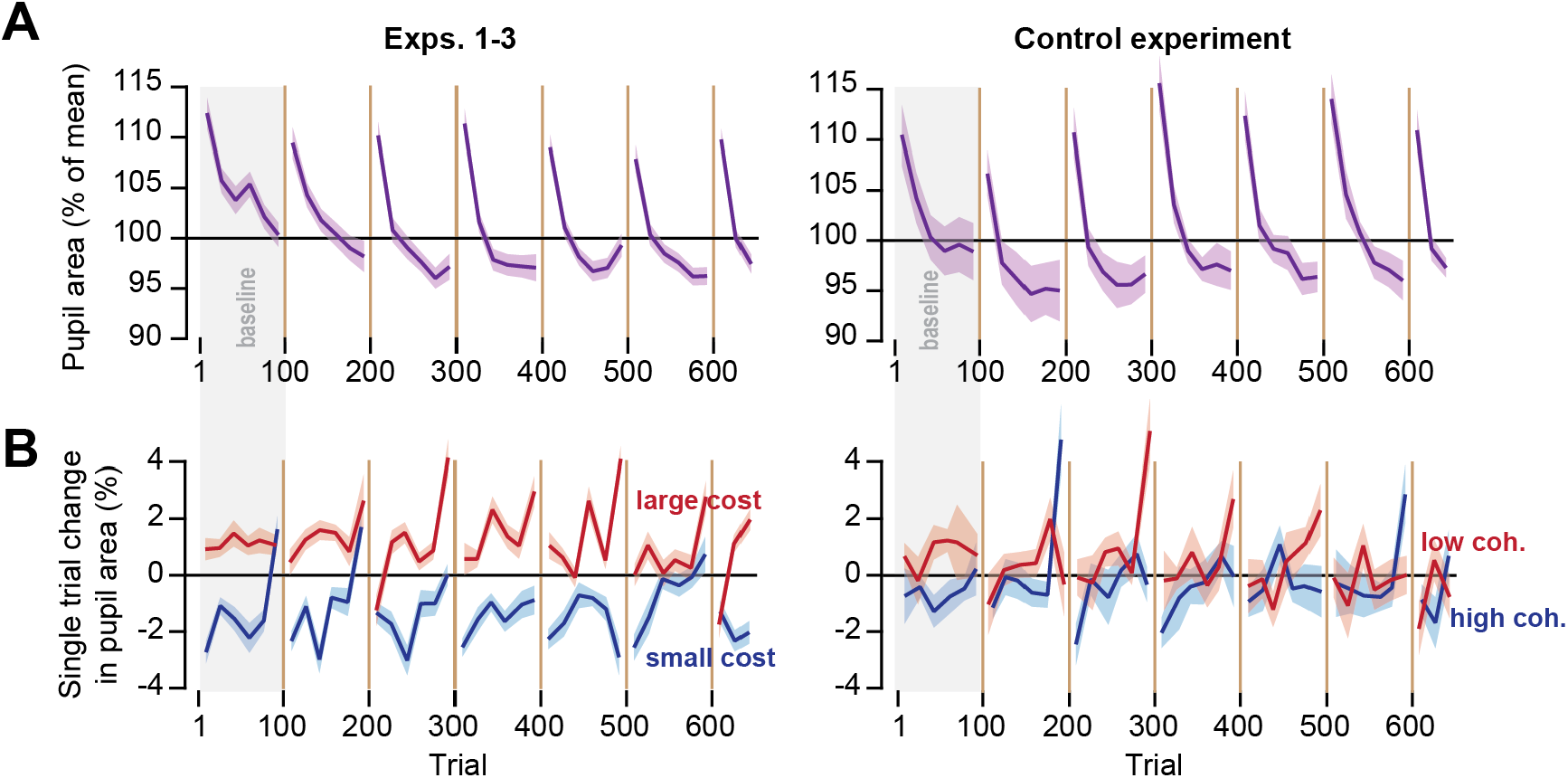
The pupil diameter changed in response to trial conditions. **(A)** Pupil area at the end of each trial, normalized to the overall mean. Pupil was most dilated when the block of trials started. It gradually constricted as the trials continued within the block, then recovered following a set break. **(B)** Effect of trial conditions on pupil diameter. The plots show the change in pupil diameter from fixation at start of a trial to onset of fixation at the start of the subsequent trial. Bin size is 8 trials. Error bars are SEM.

Next, we asked how the conditions of each trial affected pupil diameter. For each subject and each trial, we compared pupil size at center fixation (trial onset), to the fixation at the onset of the next trial before the target was displayed. This within trial response served as our proxy for how the neuromodulatory system responded to the experience of the subject during that trial.

We found that in the baseline block, the trials that were more difficult (low coherence) produced pupil dilation, whereas trials that were easy (high coherence) produced pupil constriction (Fig. 6B). The difference in the pupil response to the stimulus content of each trial was present in the baseline block of both the main group, and the control group (Fig. 6B, RM-ANOVA, trials 1-100, main effect of coherence, main experiments: F(1,59)=29.015, p<0.0005, and control experiment: F(1,19)=4.460, p=0.048). Thus, as has been noted before (Kahneman and Beatty, 1966), the difficulty of the decision-making process within each trial appeared to drive pupil dilation.

In the main experiment, as the adaptation blocks began the pupil continued to dilate in trials that were difficult and had large cost (Fig. 6B, RM-ANOVA, trials 101-650, main effect of cost, F(1,59)=51.226, p<0.0005). In the control experiment the trials were still more difficult for the low coherence stimulus, but the error cost was equalized between the two stimuli. Interestingly, in the control experiment the pupil response to trial difficulty appeared to dissipate (Fig. 6B, RM-ANOVA, trials 101-650, no main effect of coherence, F(1,19)=0.920, p=0.349). We were concerned that this difference in the two groups may have been because of the larger group size in the main experiment. However, the statistical pattern was also present in each of the main experiments (RM-ANOVA, trials 101-650, main effect of cost, Exp-1: F(1,19)=32.439, p<0.0005, Exp-2: F(1,19)=10.752, p=0.004, Exp-3: F(1,19)=7.873, p=0.011).

In summary, the pupil progressively constricted during each block of trials, suggesting a waning of attention, but then dilated following the set break, suggesting partial recovery. Within each trial the pupil diameter responded to the trial conditions: low coherence stimuli constituted more difficult trials, and in those trials the pupil dilated. However, in the adaptation block, trials in which the movement error carried a large cost continued to produce pupil dilation, as well as greater learning. In contrast, both effects diminished when we maintained trial difficulty but equalized the cost of error (control experiment).

## Discussion

When movements produce an unexpected outcome, the nervous system often produces a reflexive response that corrects for error. This corrective movement consumes time, potentially providing an implicit loss that can modulate learning from error. Here, we used saccade adaptation as a model of sensorimotor learning and explored whether imposing a cost on the time spent correcting for error could modulate learning from that error. We found that an increased error cost coincided with increased rates of adaptation. In contrast, task difficulty and reward rate altered reaction time, producing movements that responded sooner to the more rewarding stimulus. However, while reaction time was faster toward the stimulus that provided higher rate of reward, learning was greater from the stimulus that carried a larger error cost.

In a control experiment, we maintained the disparity in the rate of reward of the stimuli but equalized the error cost. This led to elimination of the effect on learning rate, suggesting that a principal factor in modulating learning from error is not the absolute rate of reward, but rather a cost that links errors with the change that they impose on probability of success.

Earlier studies have considered the effects of explicit inducements on retention of motor memories (Abe et al., 2011; Galea et al., 2015; Wachter et al., 2009). For example, Abe et al. (2011) showed that monetary reward was effective in improving retention of motor memories during a force-tracking task. Galea et al. (2015) also found that during a visuomotor rotation task presence of reward was associated with greater retention. In contrast, Steel et al. (2016) reported that during serial-reaction-time and force-tracking tasks, neither reward nor punishment benefitted retention. We implemented a spontaneous recovery paradigm and assessed how cost of error affected error sensitivity, as well as trial-to-trial retention. We found that while cost of error up-regulated sensitivity to error, it did not have a significant effect on retention.

There are other factors that influence how much the brain learns from error (Herzfeld et al., 2014; Leow et al., 2020; Marko et al., 2012; Wei and Kording, 2009). For example, Marko et al. (2012) and Hanajima et al. (2015) noted that error sensitivity was relatively high for small errors and low for large errors. Herzfeld et. al. (2014) showed that in environments where errors were likely to be consistent, subjects increased their error sensitivity. Albert et al. (2020) observed that large variability in the trial-by-trial sequence of errors tended to suppress learning from error. Conscious of these potential pitfalls, we kept the perturbation size consistent over the course of the experiment, and also controlled the statistics of the error that the subjects experienced as they made saccades. Despite this, the rate of learning was greater toward the stimulus that carried a greater error cost.

We found that increasing the cost of error rescued low adaptation, suggesting a potentially causal relationship between the cost of error and adaptation rates. Previous studies (Albert et al., 2020; Kim et al., 2019) have also found that modulating error sensitivity affected the asymptote of performance during motor learning.

In our saccade task, the pupil progressively constricted as the trials wore on within a block of trials, suggesting a decline in arousal (Mathot, 2018), but then dilated following the set break at the start of the next block, suggesting a partial recovery. The resulting saw-tooth pattern in pupil diameter was reminiscent of behavioral changes during adaptation in many other experiments: rapid adaptation that follows set breaks, and gradual adaptation that ensues with progression of trials (Chen-Harris et al., 2008; Ethier et al., 2008; Xu-Wilson et al., 2009).

Within each trial, during the baseline block the pupil dilated in response to stimuli that required greater mental effort (low coherence stimuli) and constricted in response to stimuli that required smaller effort (high coherence stimuli). During the adaptation block, when the stimuli carried an error cost the pupil continued to dilate in response to the high cost, greater mental effort stimuli. However, when the error cost was equalized in the control experiment, the dilation in response to stimuli that required greater mental effort waned. These results raise the possibility that pupil dilation is not only a correlate of attention and mental effort invested in the task, but also a correlate of learning from error.

The potential link between pupil diameter and learning from error is noteworthy because it may highlight the neural mechanism with which the brain modulates learning. Changes in pupil size are due to a band of muscles that surround the pupil, which in turn are controlled by motoneurons that reside in the Edinger-Westphal nucleus in the brainstem. Neurons in the intermediate layers of the superior colliculus project to this nucleus (May et al., 2016). As a result, weak micro-stimulation of the intermediate layers of the superior colliculus can produce a transient increase in pupil diameter that reaches its peak at around 300-500 ms (Joshi et al., 2016; Wang et al., 2012). Notably, superior colliculus neurons also project to the contralateral inferior olive (Harting, 1977), which provide climbing fibers that carry error information to Purkinje cells of the cerebellum. The climbing fiber carries information regarding the visual error following conclusion of a saccadic eye movement (Herzfeld et al., 2015; Sedaghat-Nejad et al., 2019a; Soetedjo et al., 2008), which in turn guides plasticity in Purkinje cells and affects trial-to-trial change in saccade kinematics (Herzfeld et al., 2018). Notably, the amount that the cerebellum learns from error may be related to the state of the superior colliculus: in trials in which collicular neurons respond more strongly to the visual error, there is greater trial-to-trial learning (Kojima and Soetedjo, 2018, 2017). Thus, on the one hand the superior colliculus contains the neural machinery to control pupil size, and on the other hand, it provides information to the cerebellum regarding saccade related visual errors.

Our results raise the possibility that in trials in which there is greater error cost, pupil dilation reflects an elevated state of excitability in the superior colliculus, which in turn results in a greater response to the visual error following conclusion of the primary saccade. This in turn may produce a greater probability of complex spikes in the cerebellum, and thus a larger trial-to-trial change in behavior. The coincidence of these two effects in our experiments raise the possibility that superior colliculus may play a central role in controlling cost of error during sensorimotor adaptation.

## Acknowledgements

The work was supported by grants from the National Science Foundation (CNS-1714623), the NIH (R01-NS078311, R01-NS096083), and the Office of Naval Research (N00014-15-1-2312).

## Methods

### Subjects

A total of n=80 healthy subjects (18-54 years of age, mean±SD = 24±6, 42 females) participated in our study. The procedures were approved by the Johns Hopkins School of Medicine Institutional Review Board. All subjects signed a written consent form.

### Data collection procedure

Subjects sat in front of an LED monitor (27-inch, 2560×1440 pixels, light gray background, refresh rate 144 Hz) placed at a distance of 35 cm while we measured their eye position at 1000 Hz (Eyelink 1000). Each trial began with presentation of a fixation point (a green dot, 0.5×0.5 deg) that was randomly drawn near the center of the screen: the fixation point was placed randomly in a virtual box at −1 to +1 deg along the horizontal axis, and - 1 to +1 deg along the vertical axis, where (0, 0) refers to center of the screen. After a random fixation interval of 250-750ms (uniform distribution), the fixation point was erased and a primary target (a green dot, 0.5×0.5 deg) was placed at 15 deg to the right or left along the horizontal axis.

Removal of the central fixation and presentation of the primary target served as the go signal for the primary saccade. This saccade was detected in real-time via a speed threshold of 20 deg/s, or an eye position change of 2 deg from fixation, whichever happened first. After detecting saccade onset, the primary target was erased, and a random dot image was displayed. The image was a 3×3 deg box with invisible borders containing a 0.5×0.5 deg green dot at the center and 100 0.1×0.1 deg white dots moving at 5 ^o^/s either upwards or downwards with a predefined coherence. The location of random dot image was defined based on the trial type: during baseline trials the image was centered at primary target location. During perturbation trials the image was centered at 5 deg from the primary target toward the center of the screen. During error clamp trials the image was centered at the location of the primary saccade offset.

During adaptation trials, following the completion of the primary saccade subjects produced a corrective saccade to place the random dot image on their fovea. This corrective movement carried a cost because it reduced the time available for the subject to view the image. This is because following detection of primary saccade onset, the image was displayed on thescreen, but the duration that the image was available was limited and was a key factor in our experiment design.

In the main experiments (1, 2, and 3) the image was present for only 300 ms after primary saccade offset. This was the only time available to view the image and decide on the direction of motion of the random dots. Following this 300 ms period, the image was erased, and two targets were displayed at 5 deg above and below the image. Subjects reported their perceived direction of motion by making an upward or downward saccade. Following this decision, they received feedback regarding their decision accuracy via an auditory tone: a 1000Hz (*beep*) 30 ms long sound for a correct decision, and a 500 Hz (*boop*) 30 ms long sound for an incorrect decision. At the end of this period the decision targets were removed and the center fixation point appeared at a random location near the center of the screen, in the bounding box defined above.

### Modulating cost of error

During the adaptation phase of the main experiments, although the viewing period was set to be up to 300ms from primary saccade offset, in practice, at the beginning of learning, it was around just 150 ms due to reaction time and duration of corrective saccades. Thus, by adapting the primary saccade (reducing the size of the corrective saccade), subjects would have more time to view the image, increasing the accuracy of perceiving the direction of motion. To vary the cost of error, trials consisted of two types of stimuli. For targets on one side of the screen, coherence of the random dots was low. For this image the error cost was great: the corrective saccade took precious time away from viewing the moving dots. For targets on the other side of the screen, coherence of the random dots was high. Here the error cost was small: the time consumed by the corrective saccade was irrelevant to the ability to perceive motion of the dots.

In a separate control experiment (described below), subjects received 300 ms to view the random dot image irrespective of the level of the adaptation. This served to remove the cost of error for the low and high coherence stimuli.

### Experiment design

We performed 4 experiments consisting of 3 main experiments and one control experiment. N=20 subjects participated in each experiment. The trial sequence for Exp. 1 and 2 is shown in Fig. 1E, for Exp. 3 in Fig. 2A, and for the control experiment in Fig. 4A.

All experiments started with 50 familiarization trials (no perturbations). During these trials the images appeared at the primary target location at various coherence levels to familiarize the participants with the saccadic task and motion discrimination paradigm. The collected data during the familiarization period was excluded from analysis.

After the familiarization block, the baseline block commenced. The baseline consisted of 100 trials and ended with a 30 sec set break. In this block, subjects experienced 50 low coherence trials on one side of the screen and 50 high coherence trials on the other side. The coherence side was counter-balanced between subjects. Since each subject experienced both type of stimuli (low vs high coherence), we used a within subject comparison for all statistical analysis.

Next, subjects experience 550 gain down perturbation trials (trials 101-650), during which the random dots image was displayed 5 deg away from the primary target toward the center of the screen. The consistent experience of this perturbation gradually resulted in adaptation of the primary saccades. We asked how does cost of error modulated the rate of adaptation.

All experiments included an error-clamp period. In these trials, the perturbation was removed and the image was centered at the end position of the primary saccade.

In contrast to the main experiment in which the time to view the image was reduced because of the corrective saccade, in the control experiment (Fig. 4A) the timer did not start until the end of the corrective saccade. This made it so that the time spent correcting for error did not compete with the time needed to view the random dot motion, thus equalizing the cost of error for the low and high coherence images.

### Data analysis

Eye position data were acquired using an EyeLink 1000+ system (SR Research) at 1000 Hz. Eye position data were filtered with a second-order Butterworth low-pass filter with cutoff frequency of 100 Hz. Eye velocity data in offline analysis were calculated as the derivative of the filtered position data. Saccades were identified with a speed magnitude threshold of 20^o^/sec, and minimum hold time of 10 ms at saccade end (i.e. velocity magnitude could not exceed the cutoff for a minimum 10 ms after endpoint). Corrective saccade onset and offset were detected identically to the primary saccades, using 20^o^/sec threshold on velocity magnitude. We measured change in primary saccade amplitude with respect to the average saccade amplitude in the first block in each condition. Viewing period of the image was measured based on the amount of time that the eye was inside of the random dot stimulus box from the offset of the primary saccade up to the point that the stimulus was erased and the decision targets were displayed. Decision accuracy was measured based on the number of correct decision responses divided by the total number of trials for each condition (high vs. low coherence).

Pupil area was measured by EyeLink 1000+ system (SR Reseach) and was reported in the system’s arbitrary pixel coordinate system. We blanked this data during eyeblink events to account for divergence in eye tracking. To combine and compare the pupil data across participants and experiments, we measured the percentage change for each participant by dividing the pupil data by the average pupil area over the entire recording for that subject. To control for differences in visual stimulus properties, we computed the average normalized pupil area during 200 ms window of time when participants were fixating on the start target (0.5×0.5 deg green dot). Next, we measured the change in normalized pupil area from one trial to the next to quantify how the conditions of each trial affect pupil dilation (Fig. 6B).

Statistical analyses were performed using SPSS and general linear models, with stimulus type (e.g., low or high coherence) serving as the within-subject factor. We reported results of Repeated Measure ANOVA (RM-ANOVA) with main effects of stimulus-type and trial, and stimulus-type x trial interaction.

### State-space model of learning

After the experience of a movement error, humans and other animals change their behavior on future trials. In the absence of error, adapted behavior decays over time. Here we used a state-space model (Albert and Shadmehr, 2018) to capture this process of error-based learning. Here, the internal state of an individual *x*, changes from trials *n* to *n*+1 due to learning and forgetting.

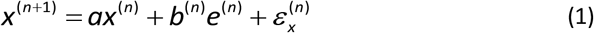

Forgetting is controlled by the trial-to-trial retention *a*. The rate of learning is controlled by the error sensitivity *b*. Learning and forgetting are stochastic processes affected by internal state noise ε_x_: a normal random variable with zero-mean and standard deviation of *σ_x_*.

While we cannot directly measure the internal state of an individual, we can measure their movements. The internal state *x* leads to a movement *y* according to:

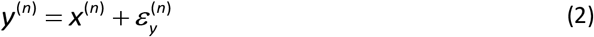

The desired movement is affected by execution noise, represented by *ε_x_*: a normal random variable with zero-mean and standard deviation of *σ_x_*. To complete the state-space model described by Eqs. 1 and 2, we must operationalize the value of an error, *e*. In sensorimotor adaptation, movement errors are determined both by motor output of the participant (*y*) and the size of the external perturbation (*r*):

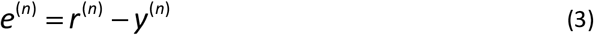

We used Eqs. (1-3) to estimate the trial-to-trial retention *a* and error sensitivity *b* during each experiment design.

---

**Supplementary Fig 1.**
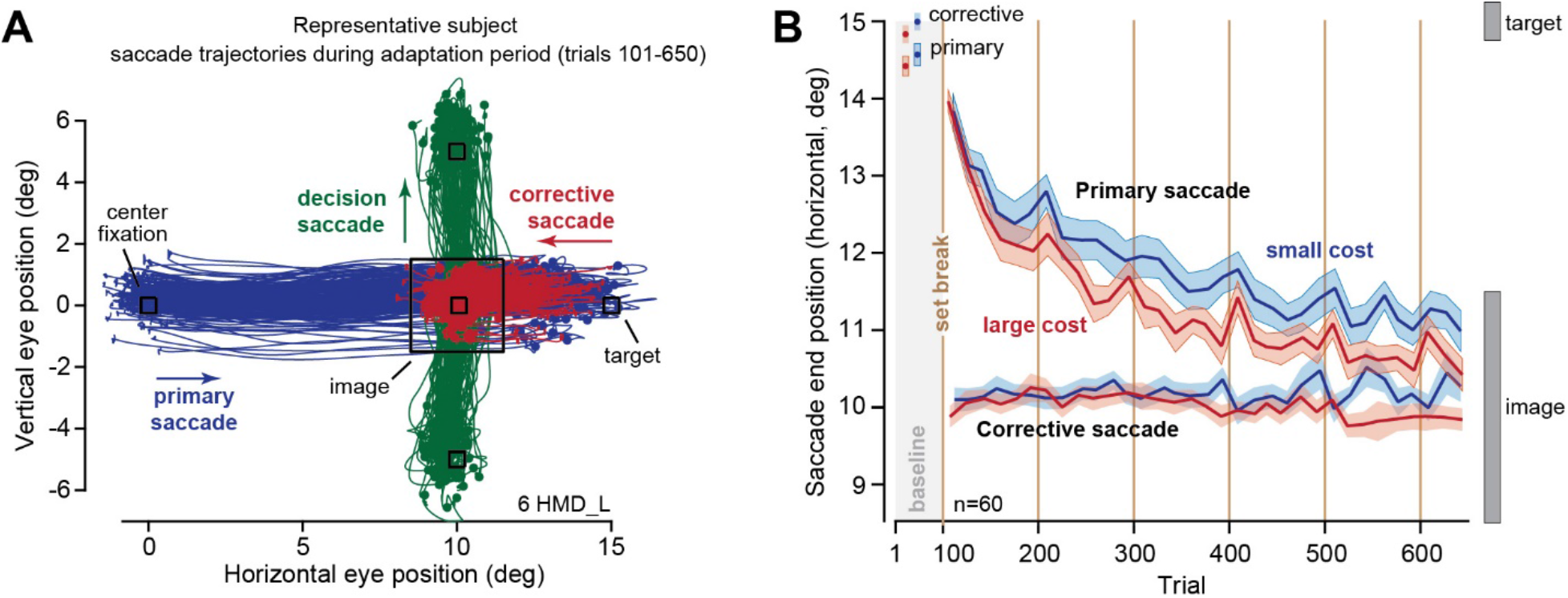
Saccade trajectory and end positions for a representative subject and for the main groups. **A**. Data for a representative subject (6_HMD_L) during the adaptation trials for the target on the right. Primary saccade is followed by a corrective saccade, and then a decision saccade. **B**. End position of the primary and corrective saccades for all subjects in the main experiments (n=60). In the baseline condition, the image was centered at the target, located at 15^o^ along the horizontal axis. Primary saccades were slightly hypometric. They were followed by a corrective saccade that brought gaze to center of the image. In the adaptation trials (101-650), the image was centered 5^o^ away from the target. The corrective saccade again brought gaze to near center of the image. This movement carried a large cost for some images, and a small cost for other images. The rate of adaptation in the primary saccade was greater when the cost of error was large. Bin size in B is 8 trials. Error bars are SEM.

**Supplementary Fig 2.**
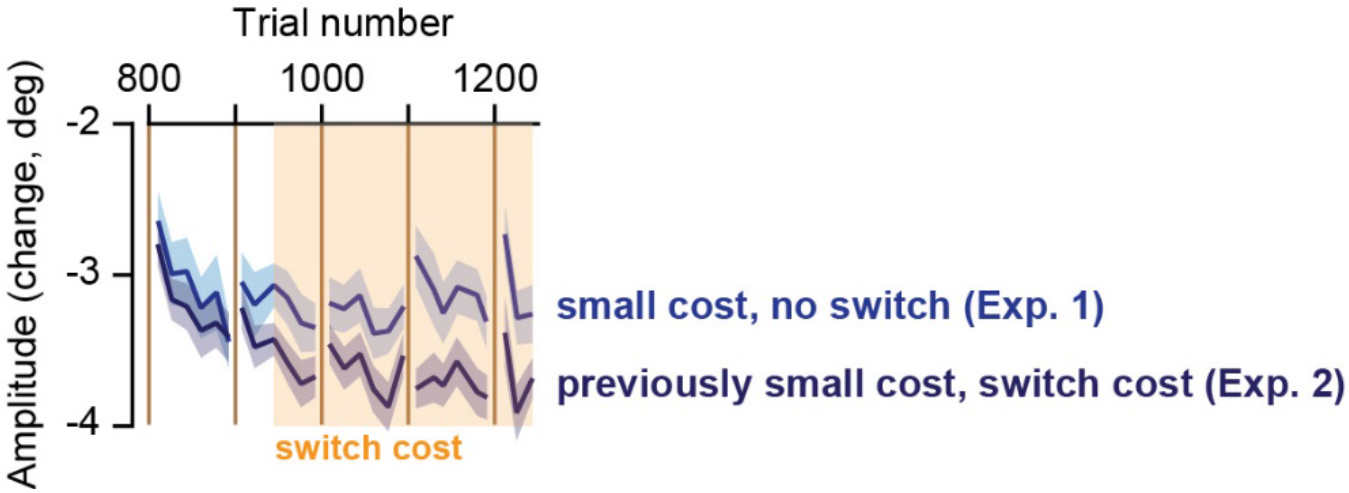
Increasing the error cost from small to large appeared to rescue adaptation levels. Data from Experiments 1&2. In Exp. 1, the side that had small cost retained that cost. In Exp. 2, the side that had small cost was suddenly switched to a large cost. Bin size is 8 trials. Error bars are SEM.

**Supplementary Fig 3.**
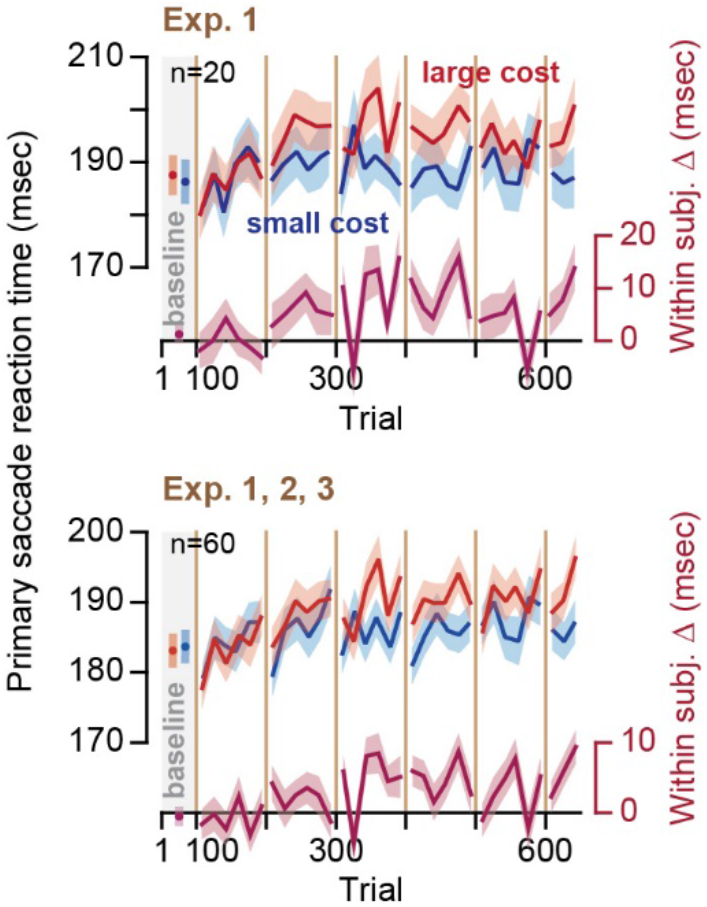
Reaction times of primary saccades were faster toward the stimuli that promised greater reward rate, not the stimuli that carried a greater cost. Note that the reaction times for the two stimuli start at approximately the same value but separate as the experiment continues. Bin size is 8 trials. Error bars are SEM.

